# *Fasciola hepatica* fatty acid binding protein (Fh12) induces apoptosis and tolerogenic properties in murine bone marrow derived dendritic cells

**DOI:** 10.1101/2021.03.29.437545

**Authors:** Caleb Ruiz-Jiménez, Daiana P. Celias, Bianca Valdés, Willy D. Ramos-Pérez, Laura Cervi, Ana M. Espino

## Abstract

In a previous study we demonstrated that *Fasciola hepatica* fatty acid binding protein (Fh12) significantly suppress macrophage function by inhibiting IL-6, IL-1B, tumor necrosis factor (TNF) and IL-12 production in TLR4-stimulated murine macrophages, an effect mediated through the signaling of CD14 co-receptor without affecting the viability of these cells. Given that dendritic cells (DCs) are immune cells that play a central role in the initiation of primary immune responses and that are the only antigen-presenting cells capable of stimulating naive T-cells, in the present study we investigated the effect of Fh12 on DCs. We found that Fh12 exerts a strong suppressive effect on activation and function of DCs. However, in contrast to the effect observed on macrophages, Fh12 induces early and late apoptosis of DCs being this phenomenon dose-dependent and CD14-coreceptor independent. At low concentration Fh12 modulates the LPS-induced DCs maturation status by suppressing the MHC-II, and co-stimulatory molecules CD40 and CD80 surface expression together with the pro-inflammatory cytokines IL-12p70 and IL-6 production whereas increase the IL-10 levels. Besides, Fh12 decreased the ability of LPS-activated DCs to induce IFNg production against allogeneic splenocytes, while increasing IL-4 production. We have described for the first time the ability of Fh12 to modify selectively the viability of DCs by apoptosis induction. The selective diminution in DCs survival could be a *F. hepatica* strategy in order to prevent a host immune response during the earliest phases of infection.

## INTRODUCTION

*Fasciola hepatica* is a helminth parasite that causes fascioliasis, a chronic disease that affects around 17 million people worldwide. Fascioliasis also leads to economic losses in livestock, estimated in more than $3 billion annually (Mas-Coma et al., 2005). The survival of helminths in the host over long periods of time is the result of a dynamic co-evolution that results in a predominant Th2/Treg immune response that is only achieved by suppressing the Th1-inflammatory response (Donnelly et al., 2008; O’Neill et al., 2001). Excretory-secretory products (ESPs) and tegumental antigens (FhTeg), which are a complex mixture of antigens, have been largely implicated as responsible for the immune modulation (Anuracpreeda et al., 2006; Brady et al., 1999; Donnelly et al., 2008). These antigens exert a strong influence on the activation status and function of the antigen-presenting cells (APCs) at the early stages of infection. In this regard, ESPs and FhTeg have shown to induce alternative activation of macrophages (Adams et al., 2014; Donnelly et al., 2005; Flynn et al., 2007) and partial activation of dendritic cells (DCs) (Hamilton et al., 2009). Moreover, ESPs have also shown to induce apoptosis of eosinophils (Serradell et al., 2007) and macrophages (Guasconi et al., 2012) during the early stages of parasite infection likely as a mechanism to prevent pro-inflammatory functions of these cells (Adam-Klages et al., 2005).

We have demonstrated that *F. hepatica* fatty acid binding protein (FABP), an antioxidant molecule with essential functions for parasite metabolism and that has been many times identified in the FhTeg and ESPs (Hacariz et al., 2012; Morphew et al., 2012; Wilson et al., 2011), possesses powerful anti-inflammatory functions. Native (Fh12) and recombinant (Fh15) variants of FABP have shown to significantly suppress the production of IL-1β and TNF from bone-marrow derived macrophages (BMDM) stimulated with LPS *in vitro* via toll-like receptor-4 (TLR4) (Martin et al., 2015; Ramos-Benitez et al., 2017). FABP exerts this effect by suppressing the expression of the CD14-coreceptor and the phosphorylation of various kinases downstream TLR4 signaling cascade (Martin et al., 2015). Moreover, Fh12 has also shown to significantly suppress a large number of pro-inflammatory cytokines in a mouse model of septic shock. Also, to impair the murine BMDM function by inhibiting their phagocytic capacity without provoking any toxic or apoptotic effect on these cells (Martin et al., 2015).

In the present study we focus on studying the effect of Fh12 on activation and functionality of DCs, which play a relevant role during the initiation of immune response in the recognition of helminth or their products and the subsequent promotion of Th2/ Treg development. In contrast to the effect observed on macrophages, we found that Fh12 modulate the activation and function of DCs by promoting the early and late apoptosis. The apoptotic effect of Fh12 on DCs was found to be dose-dependent and not mediated by the CD14-coreceptor. Concurrently, we demonstrated that F12 is capable of inducing regulatory features on LPS-activated DCs, thereby impairing their capacity to prime naïve T-cells thus, inducing a phenotype able to promote the development of anti-inflammatory responses.

## MATERIALS AND METHODS

### Animals

Six-to 8-week-old inbred female C57BL/6 and BALB/c mice were indistinctly purchased from Charles River Laboratory (Wilmington, MA) or from the Faculty of Veterinary Sciences, National University of Litoral (UNL, Argentina). B6.129S4 CD14 knockout (CD14KO) female mice (C57BL/6 background), 6–8-week-old, were purchased from Jackson Laboratory (Bar Harbor, ME). The animal studies were performed at the Animal Resources Center of the Medical Sciences Campus, University of Puerto Rico in accordance with guidelines and protocols approved by the Ethics Institutional Animal Care and Use Committee (MSC-IACUC, Protocol No. 7870215) and Faculty of Chemical Sciences, National University of Córdoba (Approval Number HCD 1637) in strict accordance with the recommendation of the Guide to the Care and Use of Experimental Animals published by the Canadian Council on Animal Care (OLAWAssurance number A5802-01).

### Fh12 Purification

Fh12 was purified from a whole-fluke extract of adult *F. hepatica* using as previously described (Espino et al., 2001). Briefly, whole fluke extract was first subjected to ultracentrifugation at 30,000 g followed by gel filtration chromatography with Sephadex G-50 (XK 26/1000 column). Fractions containing proteins in the range of 1.5-30kDa were collected, pooled and subjected to two consecutive preparative isoelectric focusing (IEF) at pH 3-10 (first separation) and pH 5-7 (second separation) using a Rotofor Cell (Bio-Rad). The individual IEF fractions were harvested, and their pH value and OD 280 nm determined. Each aliquot was subjected to SDS-PAGE and the proteins were visualized by coomasie blue. Fractions from the second IEF run that exhibited a single polypeptide band of around 12-15kDa were manually excised from the gel, washed twice with double-distilled water, digested with sequencing-grade trypsin (Promega, Madison, WI) and analyzed by MALDI and MS/MS as previously described (Morales and Espino, 2012). After confirming the presence of Fh12-FABP as a unique component, these fractions were pooled (**Fig-1S**).

**Figure-1S.**
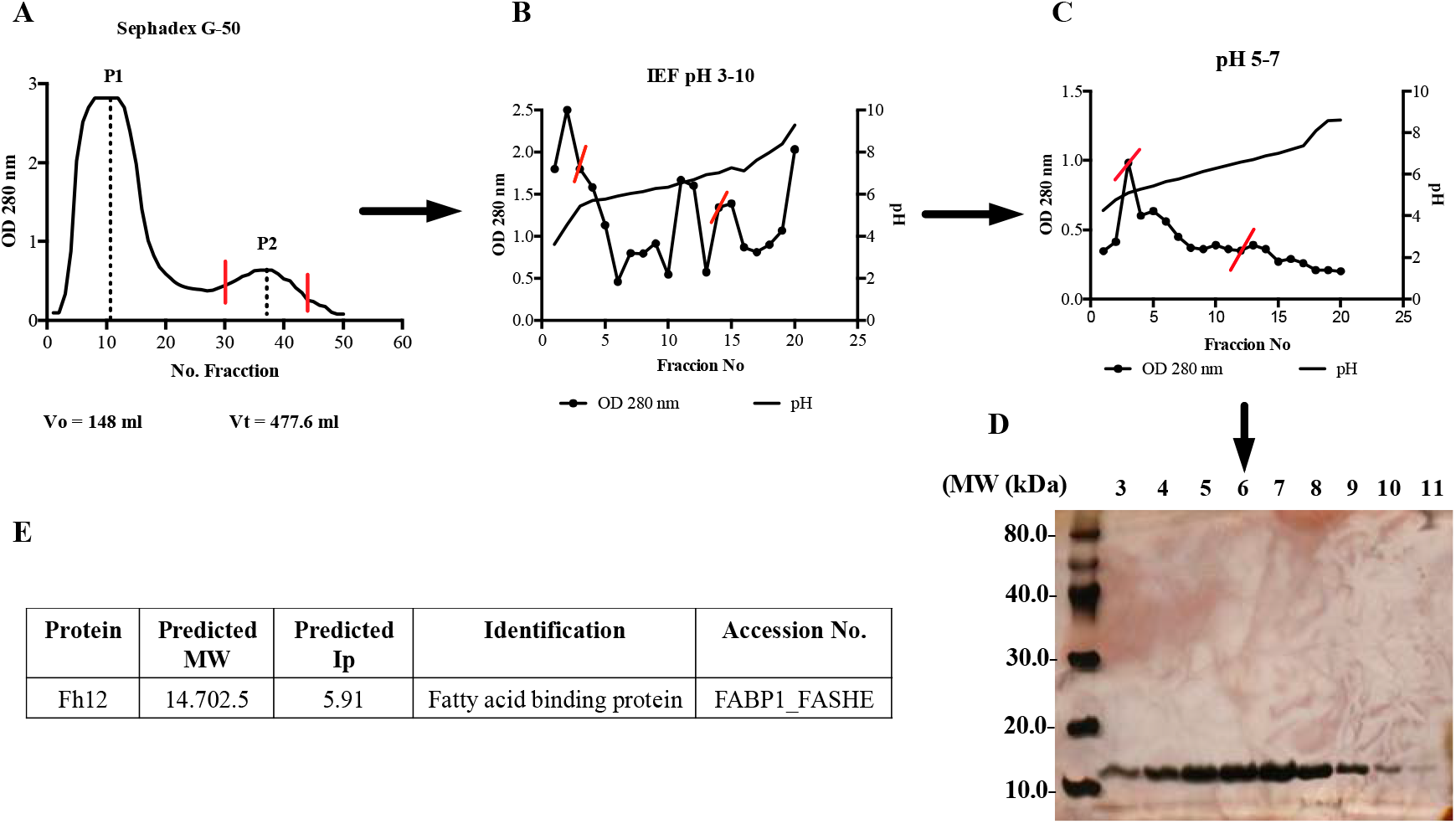
Purification of native *F. hepatica* fatty acid binding protein (Fh12). **(A)** *F. hepatica* whole worm extract was loaded onto a gel filtration chromatography with Sephadex G-50 (XK 26/100 column). Fractions eluted in peak-2 (P2) containing proteins among 1.5 to 30kDa were pooled. (**B-C**) P2 was dialyzed against 1% glycine containing 2% ampholytes pH 3-10 (first run) or pH 5-7 (second run) and then loaded onto a liquid isoelectric focusing system (Rotofor, Bio-Rad). Individual fractions were harvested and their pH and absorbance at 280nm were measured. Red lines on figures indicate fractions that were selected for pooling and subsequent purification step. **(D)** Fractions 3 to 11 from the IEF run with pH 5-7, which contain proteins with Ip between 4.61 to 5.9 were analyzed by 15% SDS-PAGE stained with silver stain to corroborate the presence of polypeptide of 12kDa. **(E)** Fractions 3 to 11 containing the 12kDa polypeptide were pooled, dialyzed against PBS, concentrated to 1mg/ml and reanalyzed by SDS-PAGE (line-1) and western-blot (line-2) using a specific rabbit polyclonal antibody against Fh12 (17). **(F)** Polypeptide band was excised from gel and analyzed by matrix-assisted laser desorption ionization 9MALDI) and tandem mass spectrometry (MS/MS). Analysis revealed the presence of fatty acid binding protein.

### Endotoxin removal

Endotoxins were removed from the Fh12 by using the polymyxin-B (PMB) column according to the manufacturer’s instructions, and the levels of endotoxins were measured using the Chromogenic *Limulus* Amebocyte Lysate QCL-1000 Assay (Lonza, Walkersville, MD). Fh12 was considered endotoxin free when endotoxin levels gave similar to background levels (<0.1 EU/ml). Protein concentration was adjusted to 1 mg/ml as determined by the bicinchoninic acid (BCA) method using a Pierce protein assay kit (Pierce, Cambridge, NJ).

### Myeloid DC generation and stimulation

DCs were generated as previously described (Falcon et al., 2010), with slight modifications. Briefly, bone marrow was collected from femurs and tibia of BALB/c mice and then seeded into bacteriological Petri dishes at 4 × 10^6^ in 10 ml of complete RPMI 1640 medium supplemented with 2mM L-glutamine (Life Technologies, Gaithersburg, MD), 100U/ml penicillin and 100μg/ml streptomycin, 10% heat-inactivated endotoxin-free fetal calf serum (Sigma-Aldrich) and 20ng/ml GM-CSF (R&D System) at 37°C, 5% CO_2_. On day 3, an additional 10 ml of medium containing 20ng/ml GM-CSF was added. At day 6, 10 ml of culture supernatant was removed and replaced with fresh culture medium containing GM-CSF. On day 8, the supernatant was removed and replaced with medium without GM-CSF, and cells were harvested 18h later (day 9). After this time, >85% of harvested cells were DCs (MHC class II^+^, CD11c^+^). For stimulation experiments, DCs were seeded into 96-wells plates at 4×10^5^ cells in complete RPMI. DCs were treated for 18h with Fh12 (2.5µg/ml or 1µg/ml LPS (*E. coli* 0111: B4; Sigma-Aldrich). In the inhibition experiments, DCs were treated for 1h with Fh12 (2.5µg/ml) and further stimulated with LPS (1µg/ml). Cells treated with PBS were used as controls in both experiments. Supernatants from cultured DCs were tested for the production of IL-10, IL-12p70 (eBioscience, USA) and IL-6 (BD Pharmingen, USA) by sandwich ELISA.

### Cell viability and Annexin-V and 7AAD assay

DCs treated with Fh12 (2.5, 5, 10 and 15 µg/ml) in the presence or absence of LPS (1µg/ml) was incubated with 50μl MTT (sodium 3′-[1-(phenylaminocarbonyl)-3,4-tetrazolium]-bis (4-methoxy-6-nitro) benzene sulfonic acid hydrate) labeling reagent (Roche Life Science, USA) to each well. The absorbance was read at 480 nm. The percentage of viable, necrotic, and apoptotic DCs after treatments were determined by flow cytometry using surface Annexin V detection and 7-Amino-actinomycin D (7-AAD) incorporation (BD Biosciences, USA). DCs were washed with PBS, and then washed twice with annexin-V binding buffer (10mM HEPES, 140 mM NaCl, 2.5 mM CaCl, pH 7.4) and resuspended in 100μl of annexin-V binding buffer before being incubated with FITC-conjugated annexin-V (0.5μg / 2 × 10^5^ cells) and 0.5μl (7-AAD) solution. After 15 min of incubation at room temperature (RT) in the dark, an extra amount of 400μl of annexin-V binding buffer was added to cells, and cells were analyzed by flow cytometry. DCs were gated on the basis of their forward and side light scatter. If there is an alteration in the membrane integrity (due to externalization of phosphatidylserine), annexin-V detects both early- and late-apoptotic cells. Thus, the simultaneous addition of 7-AAD, which does not enter healthy cells with an intact plasma membrane, discriminates between early apoptotic (annexin V-positive and 7AAD-negative), late-apoptotic (both annexin V- and 7AAD-positive), necrotic (annexin V-negative and 7AAD-positive) and live (both annexin V- and 7AAD-negative) cells (Ardestani et al., 2012).

### DCs Flow Cytometry

Expression of cell surface markers on DCs treated as described above was quantified by two-color flow cytometry using allophycocyanin-, and phycoerythrin-conjugated antibodies specific for CD80, CD40, MHC-II, and CD11c (BD BioSciences). Appropriate labeled isotype-matched antibodies were used as controls. Cell acquisition was performed using FACS Calibur equipment and MACSQuant Analyzer - Miltenyi equipment, whereas analysis of results was performed using FlowJo software (FlowJo, LLC).

### Allogeneic mixed lymphocyte reaction (MLR)

DCs from BALB/c mice were previously treated with Fh12 (2.5µg/ml), LPS (1µg/ml), or for 1h with Fh12 (2.5µg/ml) and further stimulated with LPS (1µg/ml) for 18h. Then the cells were cultured in U-bottom 96-well plates with C57BL/6 splenocytes (2×10^5^ cells/well) for 5 days at a ratio of 5:1. For cytokine determination, supernatants were collected after 48h for IFN-γ, and 72h for IL-4 and IL-13 detection.

### Statistical analysis

All experiments were repeated twice with in different days with three replicates for each determination, and equivalent results were obtained. Data are expressed as mean ± standard error of the mean (S.E.M.) and analyzed statistically using the Student *t*-test. Comparisons of the values for multiple groups were made using one-way ANOVA. Statistical analyses were made using GraphPad Prism software (Prism-6). Statistical significance was assumed at the *p*-value of <0.05.

## RESULTS

### Fh12 induce apoptosis of myeloid DCs

Given in our previous studies we had demonstrated that the treatment of murine macrophages with different concentrations of Fh12 alone or combined with LPS suppressed the capacity of these cells to express pro-inflammatory mediators without affecting the cell viability (Martin et al., 2015); in the present study, we determined whether Fh12 could exert a similar effect on DCs. For this, bone marrow derived DCs were treated for 18h with different concentrations of Fh12 or with Fh12 for 1h and further stimulated with LPS. Results of the MTT-viability test revealed that the treatment of DCs with low Fh12 concentrations (among 2.5 to 5μg/ml) alone or combined with LPS seems to maintain the viability of cells as compared with those cells treated only with PBS or LPS alone. However, at Fh12 concentrations of 10 and 15μg/ml, the absorbance values significantly dropped, which indicates that at these Fh12 concentrations, the viability of cells was seriously compromised (*p*<0.0001) (**Fig. 1**). To determine whether Fh12 could be inducing apoptosis, Fh12-treated DCs were stained with annexin-V and 7AAD and analyzed by flow cytometry. Results demonstrated that ∼91% of cells remain viable when are treated with medium alone for 18h. However, when cells are treated with Fh12 the viability of cells drop significantly. At Fh12 concentrations of 2.5 or 5μg/ml ∼ 52% and 41.9% of DCs remained viable, whereas that 46.5% (23.6% early apoptosis and 22.9% late apoptosis) and 52.9% (18.8% early apoptosis and 34.1% late apoptosis) was in apoptosis, respectively. The number of apoptotic cells increased significantly by 80.7% (22.1% early apoptosis and 58.6% late apoptosis) and 88.7% (18.3% early apoptosis and 70.4% late apoptosis) when the Fh12 concentration increased to 10μg/ml and 15μg/ml, respectively (**Fig. 2**). These results clearly demonstrate that Fh12 promotes the late apoptosis of DCs in a dose-dependent manner.

**Figure-1.**
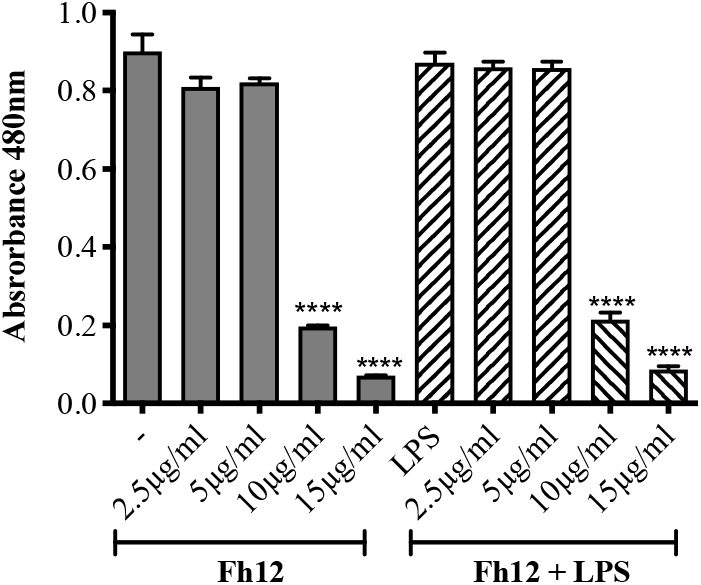
MTT-viability assay of Fh12 treated DCs. Murine DCs were seeded into 96-wells plates at 4×10^5^ cells in complete RPMI plus 20ng/ml GM-CSF and then treated with Fh12 (2.5, 5, 10, 15µg/ml) in the presence or absence of 1µg/ml LPS (*E. coli* 0111:B4). Control cells were treated with Fh12 or LPS alone. After 18h of incubation at 37°C, 5% CO_2_ 50μl MTT (sodium 3′-[1-(phenylaminocarbonyl)-3,4-tetrazolium]-bis (4-methoxy-6-nitro) benzene sulfonic acid hydrate) labeling reagent was added to each well and the absorbance was read at 480 nm. Viability of cells significantly dropped (**** *p*<0.0001) at Fh12 concentrations ≥10μg/ml.

**Figure-2.**
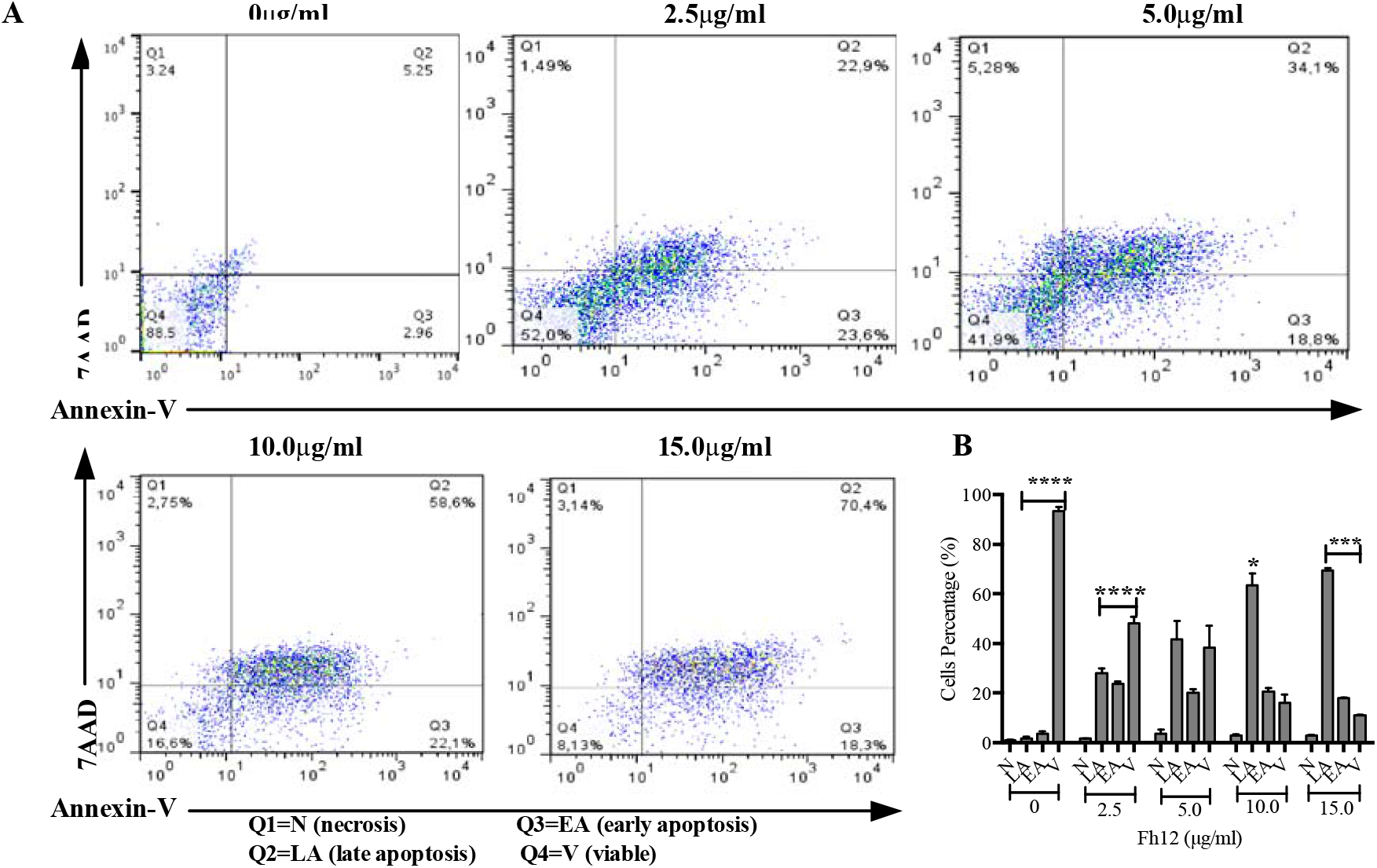
Fh12 induces apoptosis of DCs, as measured by Annexin-V binding to externalized phosphatidylserine. Myeloid DCs from C57BL/6 mice were treated with Fh12 (2.5, 5, 10, and 15 μg/ml) or remain untreated for 18 hours. After incubation, the cells were harvested and analyzed by two-color flow cytometry for Annexin V and 7-AAD. Cells were gated based on the CD11c expression. **(A)** Histogram representative of an experiment showing that the apoptosis of DCs induced by Fh12 is dose-dependent. **(B)** Average percentages and their SEM of at least three independent experiments showing that the percentage of DCs viable (V) at Fh12 concentration of 2.5μg/ml is significantly high (****p<0.0001) compared to the percentage of cells in early (EA) or late apoptosis (LA). In contrast, the percentage of DCs in apoptosis significantly increased at Fh12 concentration of 10μg/ml. (\**p*= 0.0146) and 15μg/ml (\*\*\**p*<0.0008).

### Fh12 induces apoptosis of DCs in absence of CD14 co-receptor

Since in our previous studies with murine macrophages we had demonstrated that Fh12 targeted the CD14 coreceptor as a mechanism to suppress the expression of pro-inflammatory cytokines and prevent the phagocytic capacity of macrophages (Martin et al., 2015), and given that CD14 has been involved in the regulation of programmed cell death (apoptosis) in immune and no-immune cells (Frey and Finlay, 1998; Heidenreich et al., 1997), we investigate whether Fh12 could require CD14 to induce apoptosis of DCs. Myeloid DCs were collected from CD14 KO mice and fully differentiated *in vitro*. Further, cells were cultured and treated with 15μg/ml Fh12. Results demonstrated that in the absence of CD14 Fh12 induces similar apoptosis levels that those observed in DCs of wild type animals (more than 70% **Fig. 3**). This result indicates that CD14 signaling is not involved in the apoptosis mechanism induced by Fh12.

**Figure-3.**
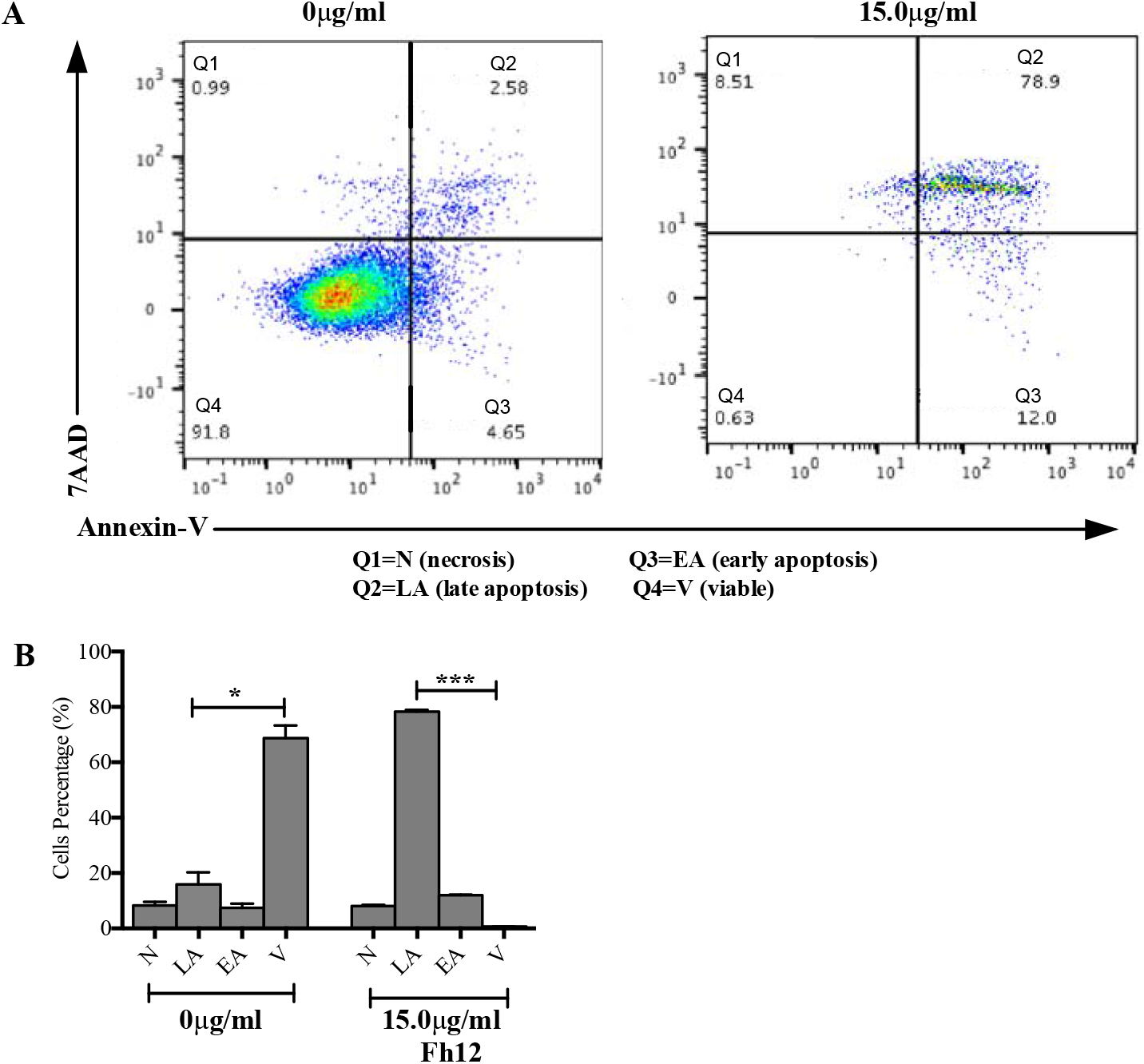
Fh12 induces apoptosis of DCs in absence of CD14 coreceptor. Myeloid DCs from CD14 KO C57BL/6 mice were treated with Fh12 (15μg/ml) for 18 hours or remain untreated. After incubation, the cells were harvested and analyzed by two-color flow cytometry for Annexin V and 7-AAD. Cells were gated based on the CD11c expression. **(A)** Histogram representative of an experiment showing the apoptosis of DCs induced by Fh12. **(B)** Average percentages and their SEM of at least three independent experiments showing that the percentage of DCs viable (V) compared to apoptotic (EA or LA) or necrotic cells (N) in untreated CD14 KO cells is significantly high (\**p*=0.144) whereas the number of DCs in late apoptosis (LA) compared to viable (V) cells is significantly high (\*\*\*\**p*<0.0001).

### Fh12 did not induce DCs classical maturation but affected LPS-induced DCs activation

Having observed that after treatment with low Fh12 concentrations (2.5μg/ml) near 52% of DCs still remain viable, we investigated whether at this experimental conditions Fh12 could modify the activation status of LPS-activated DCs. The cytokines and the expression of surface proteins involved in the T-cell activation and polarization were analyzed in DCs cultured with medium alone, Fh12 (2.5μg/ml), LPS (1μg/ml) or for 1h with Fh12 and further stimulated with LPS. As expected, DCs stimulated with LPS produced significantly more IL-12p70 (p=0.0006) and IL-6 (p=0.0003) than untreated DCs and did not produce IL-10. However, although Fh12 alone did not induce the production of IL-12p70, IL-6, or IL-10, it significantly suppressed the LPS-induced IL-12p70 (p<0.031) and IL-6 (p<0.0458) production. In addition, in the presence of Fh12, LPS-stimulated cells produced significantly more IL-10 (p=0.0039) than untreated cells (**Fig. 4**). Moreover, the expression of the MHC-II and co-stimulatory molecules CD40 and CD80 induced by LPS-stimulation was found significantly downregulated by the Fh12 treatment (*p*=0.0072, *p*=0.0023, and *p*=0.0398, respectively) (**Fig. 5**). Together, our data showed that Fh12 was not able to induce classical DC maturation but exerted control over the DC activation induced by LPS. These not only by virtue of its ability to inhibit expression of co-stimulatory molecules essential for T-cell activation, but also by directly suppressing the secretion of cytokines critical for the Th1 cell polarization.

**Figure-4.**
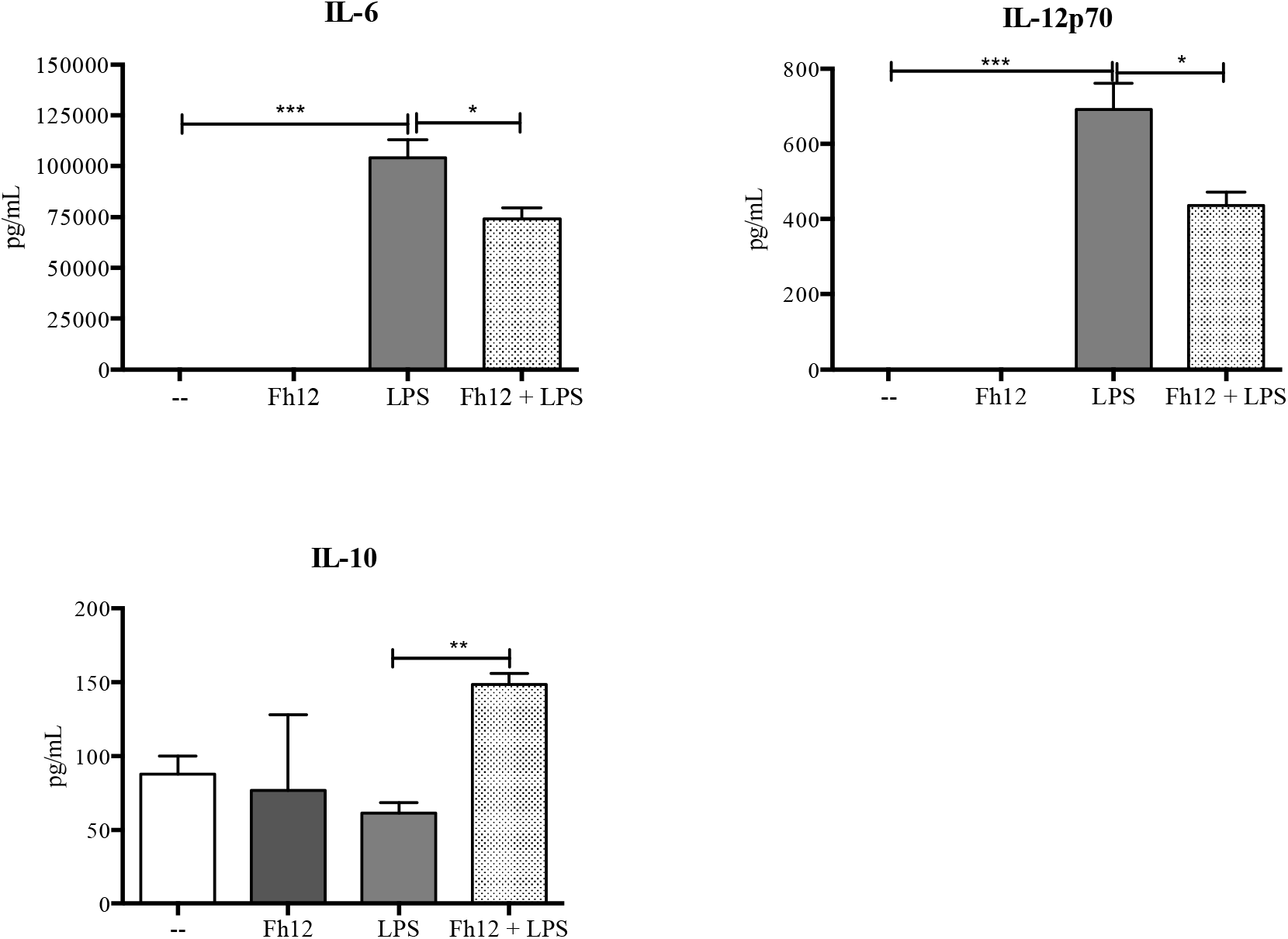
Levels of IL-6, IL-12p70, and IL-10 cytokines measured by ELISA in the supernatant of DCs treated with Fh12. DCs collected from naïve BALB/c mice were cultured and treated with medium or Fh12 (2.5μg/ml) 1h before stimulation with LPS (1μg) for 18h LPS significantly produced high levels of IL-6 (***p=0.0003), which were significantly suppressed by Fh12 (\*\*\**p*<0.0458). LPS also induced the production of IL-12p70 (***p=0.0006), which also were significantly suppressed by Fh12 (\**p*<0.0315). Although Fh12 or LPS alone failed in inducing IL-10, in the presence LPS, Fh12 significantly increases the production of IL-10 (**p=0.0039). Data represent averages + S.E.M. of at least 3 replicated from three different experiments.

**Figure-5.**
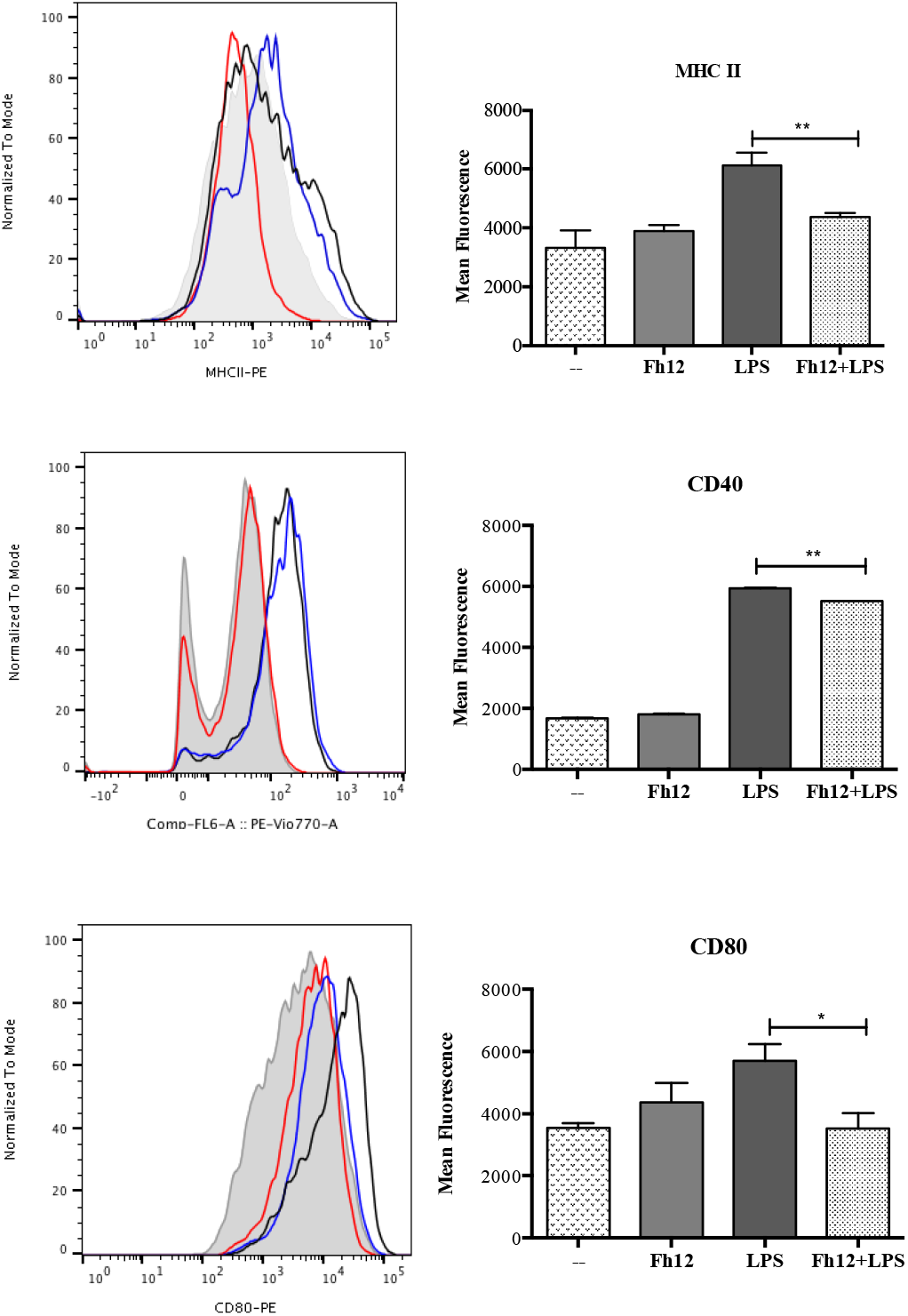
Fh12 inhibits MHC II and co-stimulatory molecules expression on LPS-activated DCs. DCs from BALB/c mice were treated for 1h with Fh12 (2.5 μg/ml) and further stimulated with LPS (1μg) for 18h. After incubation, the cells were harvested and analyzed by two-color flow cytometry for MHC-II, CD40 and CD80. Cells were gated based on the expression of CD11c. Histograms (left panel) show LPS-stimulated cells (black line), Fh12-treated cells (red line), Fh12+LPS-stimulated cells (blue line) and untreated cells (--) (gray shaded histogram). Right panel represents mean fluorescence intensity of untreated DCs or treated with Fh12, LPS or Fh12/LPS, for MHC-II, CD40 and CD80, respectively (\*\**p*=0.0072, \*\**p=* 0.0023, \**p*=0.0398).

### Fh12-treated DCs showed reduced ability to induce allogeneic responses

Given our data showing that Fh12 inhibits the maturation of LPS-treated DCs, we reasoned that the ability of these cells to prime allospecific T-cell responses could be impaired. Then, we wanted to determine which T helper profile was promoted by Fh12-treated DCs in a mixed lymphocyte reaction. To this end, immature or LPS-treated DCs or treated for 1h with Fh12 and further stimulated with LPS from BALB/c mice were then cultured with allogeneic splenocytes from C57BL6. Cultures with untreated DCs were used as control. As expected, DCs stimulated with LPS and co-cultured with allogeneic splenocytes induced the secretion of significantly higher amounts of IFN-γ (p=0.0003) compared to untreated cells and failed to induce allogeneic IL-4 or IL-13 production. In contrast, Fh12-treated DCs induced significant amounts of IL-4 (*p*=0.0022) and IL-13 (*p*<0.0001) compared to untreated cells. Importantly, LPS-stimulated DCs that were pre-treated with Fh12 and then co-cultured with allogeneic splenocytes lack the ability to induce INF-γ production (p=0.0003) whereas significantly augmented their capacity to produce IL-4 (p=0.0073). Fh12 also showed a tendency to increase the production of IL-13 in DCs stimulated with LPS and then co-cultured with splenocytes, but these increases were not found significant **(Fig. 6)**. These results suggest that Fh12 could be able to promote the activation of T-helper type-2 (Th2) responses of immature or LPS-mature DCs, whereas concurrently suppress the ability of these cells to promote the Th1-response.

**Figure-6.**
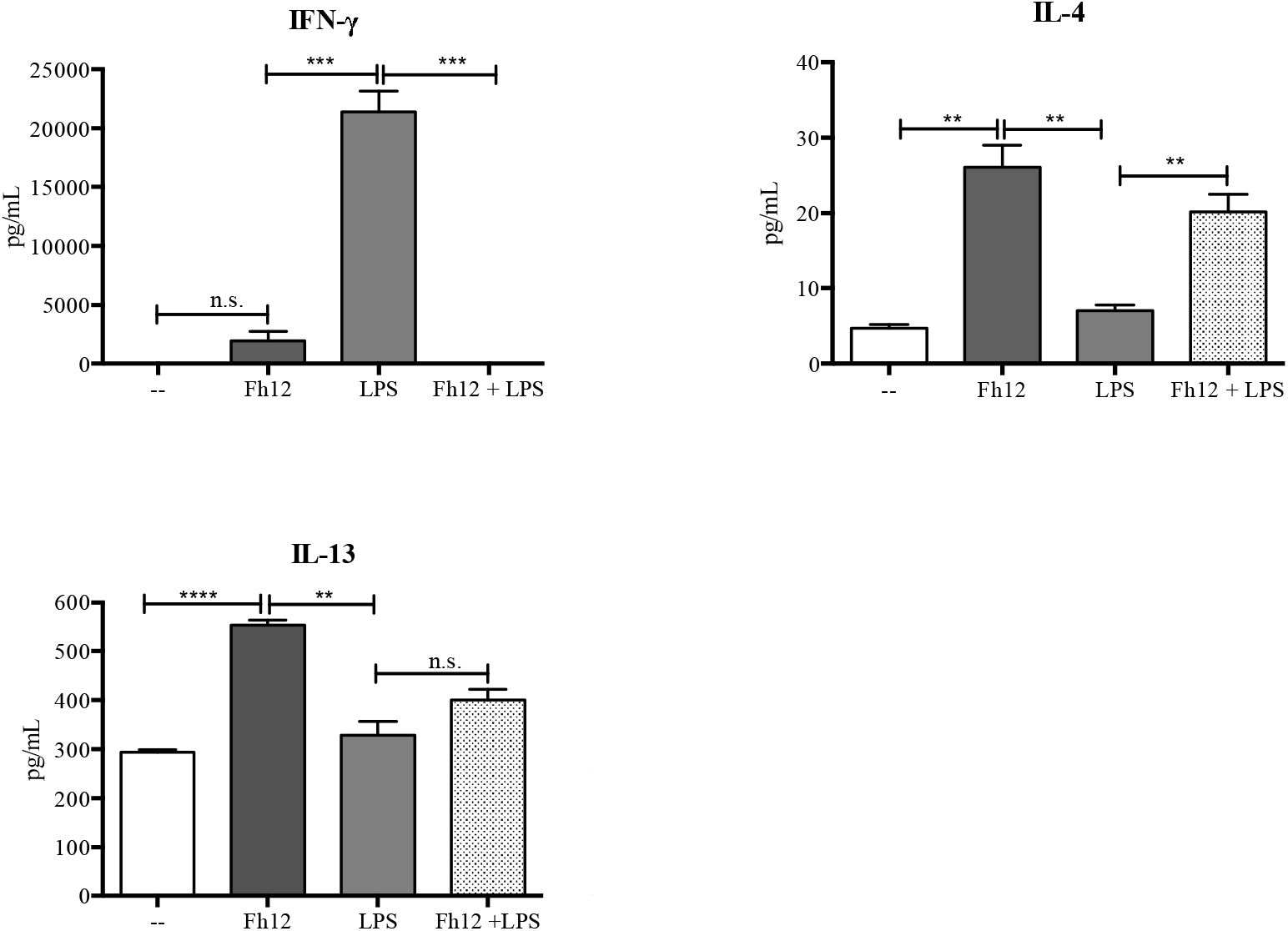
Fh12 modulates DCs capacity to prime allogeneic responses. DCs collected from BALB/c mice were treated for 1h with Fh12 (2.5 μg/ml) and further stimulated with LPS (1μg/ml) for 18h. Cells were then washed and co-cultured with CD4+ T-cells from naïve C57BL/6 mice for 5 days. Fh12 significantly suppressed the LPS-induced IFN-γ levels (\*\*\**p*=0.0003). Fh12 significantly increased the levels of IL-4 either when it was added to the culture alone (*p*=0.0022) or before the LPS-stimulation (\*\**p*=0.0073). Fh12 alone significantly increased the production of IL-13 compared to untreated cells (p<0.0001) or stimulated with LPS (*p*=0.0017). However, in the presence of Fh12 + LPS, although it was observed a trend to increase the levels of IL-13, this increase was not significant. The data represent averages with their SEM of at least 3 replicates from three independent experiments.

## DISCUSSION

*F. hepatica* can be considered a successful parasite because of its ability to migrate through the host tissues without suffering damage to finally allocate in the bile ducts, where it can survive for years. To achieve this survival, *F. hepatica* has co-evolved with the host and is able to induce a type of response that is not harmful to it. During its migration, the parasite excretes and/or secretes many products capable of modulating the immune response (Molina-Hernandez et al., 2015). Soon after the infection, the juvenile stage of *F. hepatica* crosses the intestinal wall and reaches the peritoneum inducing the recruitment of alternatively activated macrophages (Adams et al., 2014; Donnelly et al., 2005; Donnelly et al., 2008). Different reports have shown the ability of various products such as ESPs (Guasconi et al., 2011), and FhTeg (Hacariz et al., 2011) and different enzymes such as thioredoxin peroxidase (Donnelly et al., 2005), 2-Cys peroxiredoxin (Donnelly et al., 2008), fatty acid binding protein (Fh12) (Figueroa-Santiago, 2014) and more recently heme-oxygenase-1 (Carasi et al., 2017) to modulate macrophage activation toward an alternative profile. This phenotype promotes the secretion of anti-inflammatory factors, enhancing the differentiation of Th2 and Treg cells (Kreider et al., 2007). In addition to the modulation of macrophages activation, *F. hepatica* derived products such as ESPs (Falcon et al., 2010), FhTeg (Hamilton et al., 2009) and secreted proteins such as Kunitz type molecule (KTM) (Falcon et al., 2014) has been related to a down-modulation in DCs activation. It is reasonable to assume that distinct molecules, secreted or expressed by the parasite in its tegument, may participate during its migration in the prevention of an appropriate activation of DCs, contributing to an anti-inflammatory control.

Given that Fh12 is an immunogenic protein that plays an important role in nutrient acquisition and survival of the parasite and is present in ESPs and FhTeg, its interaction with DCs might be relevant. In this work, we investigate the effect of Fh12 on maturation and function of DCs. Unexpectedly, our results showed that Fh12 induces the early and late apoptosis of DCs in a dose dependent manner. Additionally, Fh12 inhibited the ability of DCs to mature in the presence of LPS. Considering that Fh12 inhibits inflammatory cytokines in LPS matured macrophages through a mechanism that involves CD14, we hypothesize that this molecule might be involved in the apoptosis. However, our results are against this hypothesis since DCs from CD14 deficient mice in the presence of Fh12 showed similar apoptosis levels of those observed in DCs from normal animals, suggesting the independence of this phenomenon from CD14.

Being a CD14 a coreceptor that participates along with the toll-like receptor-4 (TLR4) and MD-2 in the recognition of bacterial LPS (Dowling, 2018; Kitchens, 2000) its importance in the recognition of Fh12 by DCs seems to be relative. In contrast, it has been shown that CD14 is involved in the apoptotic death of LPS-stimulated DCs via NFAT activation has been demonstrated (Zanoni et al., 2011). However, unlike what happens with DCs, macrophages do not die after LPS activation. The authors argue that after activation by LPS, significant differences are generated in the signal transduction pathways in DCs and macrophages. These last cells were unable to mobilize Ca, a crucial event to induce apoptosis. Interestingly tissue-resident macrophage survival after activation is a crucial event for inflammation resolution (Zanoni et al., 2009).

Interestingly, in previous reports, the induction of apoptosis in eosinophils by ES products of *F. hepatica* through the induction of reactive oxygen species (ROS) has been shown (Serradell et al., 2007). Since Fh12 is part of the *F. hepatica* ESPs, a similar mechanism of apoptosis induced by Fh12 in DCs might be occurring. Experiments using ROS inhibitors or other apoptosis mediators should be used to dissect the mechanism by which Fh12 exerts this effect on DCs.

Based on our results showing that the lowest concentration of Fh12 induce approximately 47% of the cells in apoptosis, while low concentration of LPS (100ng/ml) induced only 17.5% of the apoptotic cells (**Fig. 2S**) and both stimuli induce different activation status in DCs, we hypothesize that the apoptosis and the modulation of DCs activation might be independent events. Data supporting this notion show that the signaling involved in survival and LPS-induced activation in DCs would be different, dependent on ERK and NF-κB, respectively (Rescigno et al., 1998). Both phenomena could occur simultaneously, and it would be difficult to determine whether they are dependent or independent events. During the allogenic responses, a mismatch between the MHC-II molecules presents in the DCs and the TCR on T lymphocytes promotes the proliferation and production of cytokines, mainly IFN-γ. This phenomenon is exacerbated when the microenvironment generated during DCs stimulation is inflammatory (Herrera et al., 2004). Our results indicate that Fh12 exerts a negative regulation on DCs maturation induced by LPS inhibiting MHCII and co-stimulatory expression generating defective activated DCs with low production of proinflammatory molecules or cytokines. The poor maturation status of DCs might explain the impaired ability of these cells to induce allogeneic responses. This effect exerted by Fh12 does not appear to be redundant since other molecules derived from *F. hepatica* such as cathepsin L1 or glutathione S-transferase inhibit IL-23 secretion from LPS-matured DCs, with the consequence of a reduction in the inflammatory Th17 response. Taken together, these results suggest that different antigens derived from *F. hepatica* migtht attenuate the DCs activation, which in turn is responsible for the decrease of both type Th1 and Th17 inflammatory responses, what otherwise it would be harmful to the parasite. The control of DCs activation, together with the apoptosis of these cells, might be useful to reduce the impact of bacterial ligands translocation during the larvae passage through the intestinal wall. Thus, our data suggest an immunomodulatory role for Fh12 on DCs function and its possible involvement in immunoevasion mechanisms.

**Figure-2S.**
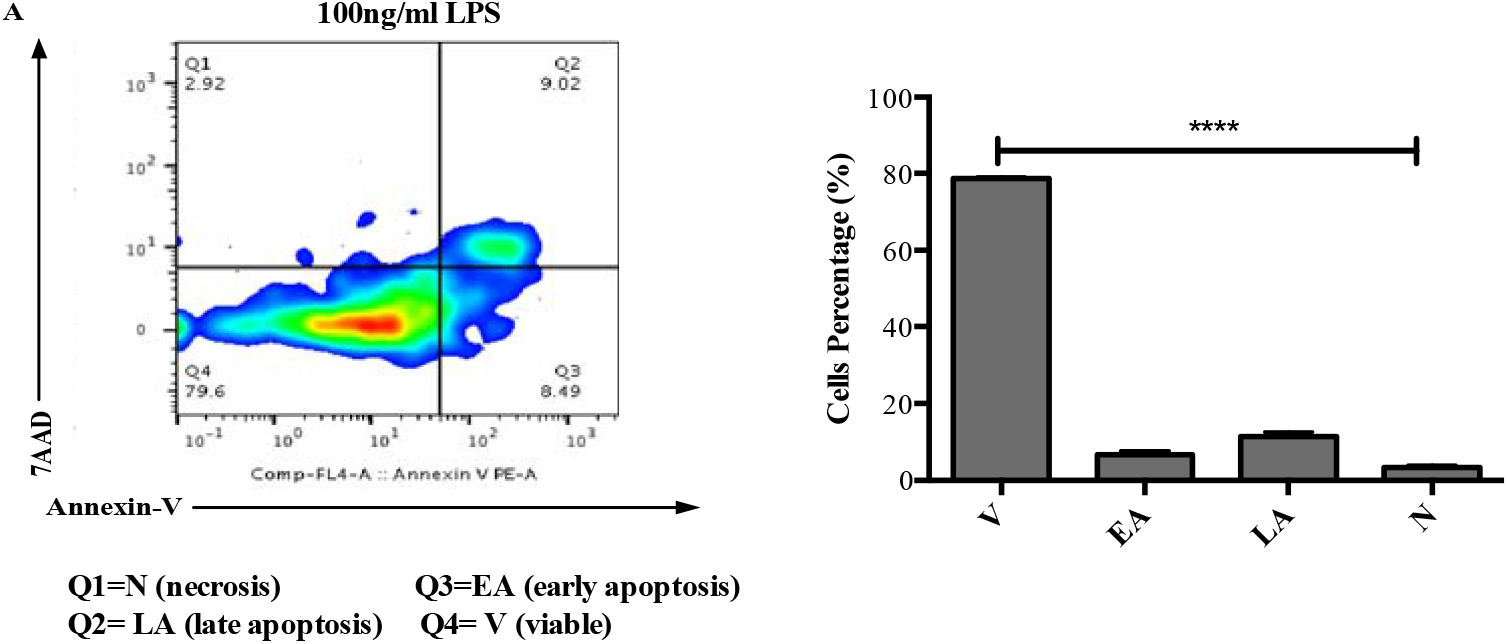
LPS does not promote significant apoptosis to DCs. Myeloid DCs from C57BL/6 mice were treated with LPS (100ng/ml) or remain untreated for 18 hours. After incubation, the cells were harvested and analyzed by two-color flow cytometry for Annexin V and 7-AAD. Cells were gated based on the CD11c expression. **(A)** Histogram representative of an experiment showing that at this experimental conditions most of cells (79.6%) remain viable and that only the 17% of cells are apoptotic. **(B)** Average percentages and their SEM of at least three independent experiments showing that the percentage of DCs viable (V) at LPS concentration of 100ng/ml is significantly high (****p<0.0001) compared to the percentage of cells in early (EA) or late apoptosis (LA).

## ACKNOWLEDGMENTS

This work was supported by a grant from the National Institute of Allergy and Infectious Diseases (NIAID) 1SC1AI155439-01 and grants from the National Institutes on Minority Health Disparities G12MD007600, R25GM061838 and 5R25GM061151. The contents of this article are solely the responsibility of the authors and do not necessarily represent the official views of the National Institutes of Health.

